# Migrating lung monocytes internalize and inhibit growth of *Aspergillus fumigatus* conidia

**DOI:** 10.1101/2020.10.12.335786

**Authors:** Natalia Schiefermeier-Mach, Thomas Haller, Stephan Geley, Susanne Perkhofer

**Affiliations:** FH Gesundheit / Health University of Applied Sciences Tyrol, Innrain 98, Innsbruck 6020, Austria; Institute of Physiology, Medical University of Innsbruck, Innrain 52, 6020 Innsbruck, Austria; Institute of Pathophysiology, Medical University of Innsbruck, Innrain 52, 6020 Innsbruck, Austria

**Keywords:** migrating lung monocytes, patrolling monocytes, non-classical monocytes, alveolar cells, fungal conidia, *Aspergillus fumigatus*

## Abstract

Monocytes are important players to combat ubiquitously present fungus *Aspergillus fumigatus*. Recruitment of monocytes to sites of fungal infection was shown *in vivo*, and purified murine and human blood monocytes are able to induce inflammatory and fungicidal mediators as well as the host cell and the fungal transcriptional responses upon exposure to *A.fumigatus*. Mononuclear tissue phagocytes are phenotypically and functionally different from those circulating in the blood and their role in antifungal defences is much less understood.

In this study, we identified a population of migrating *CD43*+ monocytes in cells isolated from rat distal lungs. These cells phenotypically different from alveolar macrophages, showed clearly distinct locomotory behaviour on the surface of primary alveolar cells resembling previously described endothelial patrolling. The *CD43*+ monocytes internalized live *A.fumigatus* conidia resulting in inhibition of conidial germination and hyphal growth. Thus, migrating lung monocytes might play an important role in local defence against pulmonary pathogens.

## Introduction

The large surface of the respiratory tract is constantly exposed to external environmental factors and elaborate cleaning systems operate to control pathogenic hazards ubiquitously present in inhaled air. The lung is populated with an intricate network of immune cells, including mononuclear phagocytes (MNPs) ^1–4^. Previous studies have reported that MNPs residing in peripheral tissues are phenotypically and functionally different to circulating monocytes. In particular, isolated lung immune cells provided evidence of several types of MNPs: monocytes, dendritic cells (DCs), resident alveolar macrophages (AMs) and interstitial macrophages ^1,5,6^. Whereas dendritic cells and macrophages are well described in terms of ontogeny, function, expression of phenotypical markers and migration ^7–11^, little is known about lung specific monocytes. Recently described “patrolling” behaviour of migrating lung monocytes as well as their localization at the interface between the capillaries and the alveoli suggest immune surveillance function of these cells ^12,13^.

Monocytes are increasingly recognized as important players against the filamentous and ubiquitously present fungus *Aspergillus fumigatus*. Two classes of circulating monocytes are found in peripheral blood and distinct tissues, like the lung ^12,14–16^: “classical monocytes” and “non-classical monocytes *(CD14^+/-(dim)^CD16^++^ in human, Ly6C^lo^CD43^+^CD62L^-^CCR2^-^ in mice, CD43^++^CD62L^-^CCR2^-^ in rats)*” and some studies define another activated phenotype, the “intermediate monocytes” *(CD14^+^CD16^+^ in human, Ly6C^int^CD43^+^CD62L^-^CCR2^-^ in mice)*. They may in fact constitute a third subset and are so far found in men and mice, but not in rats ^17–20^. The relative contribution of non-classical monocytes to the initiation of immunity^21–24^ and/or maintenance of tolerance ^25–27^ implies a dual role that is of crucial interest.

Inhaled conidia of*A.fumigatus* may overcome the upper respiratory tract defence mechanisms and reach the pulmonary alveoli, where macrophages and neutrophils are established as the keystones of host defence ^28,29^. Alveolar macrophages were shown to digest and kill conidia, while recruited neutrophils attack *Aspergillus* hyphae that occasionally escaped macrophage killing ^30,31^. Recruitment of monocytes to sites of fungal infection has also been shown *in vivo* ^32^, however the exact role of monocyte subsets in fungal killing remains unclear. Recent report by Espinosa, Jhingra et al. has shown that depletion of classical *CCR2*+ monocytes and their derivative DCs in knockout mice decreases *A.fumigatus* conidial containment and results in reduction of neutrophil conidiacidal activity ^33^. *In vitro* experiments using purified murine and human blood monocytes suggested that an *A.fumigatus* infection results in induction of inflammatory and fungicidal mediators ^30,34^ as well as the host cell and the fungal transcriptional responses ^35^. Upon recognition of fungal β-D-glucan residues by Dectin-1, conidia are phagocytosed by monocytes resulting in inhibition of conidia growth. Monocytes were also shown to secret TNF- α and iNOS in response to *A.fumigatus* ^33,36^.

In this study, we identified non-classical *CD43*+ patrolling monocytes in a primary cell mix isolated from rat lungs. We further investigated the function and migration capacity of lung monocytes in an *A.fumigatus* infection-like setting. Our results suggest that monocytes residing in distal lung tissue have a capacity to i) migrate on the surface of alveolar cells; ii) internalize and transport *A.fumigatus* conidia and iii) inhibit conidial germination and hyphal growth.

## Results

### Cells isolated form alveoli contain non-classical migrating monocytes

We have modified a previously published method ^37^ of the primary alveolar type II (ATII) cells isolation from rat lungs to include also MNPs present in the tissue (**Figure 1A-D**). Cells allowed to grow 48h on glass coverslips contained all cell types characteristic for alveolar tissue, including predominantly ATII cells, but also ATI cells, fibroblasts and MNPs. Isolated lung MNPs contained *CD45*+ cells as well as a *CD11b*+ fraction that may include AMs, DCs and monocytes (Figure 1A). CD43 was previously shown to be a specific marker of non-classical monocytes in rats ^38,39^. We observed a small portion of *CD43*+ rat monocytes (4.3% of all isolated cells, Figure 1A, n=360, see Materials and Methods). The morphology of *CD43*+ cells was strikingly different from the other cells in the mix and was characterized by a smaller cell size, elongated compact nucleus and a more polarized cell shape (**Figure 1A**).

**Figure 1.**
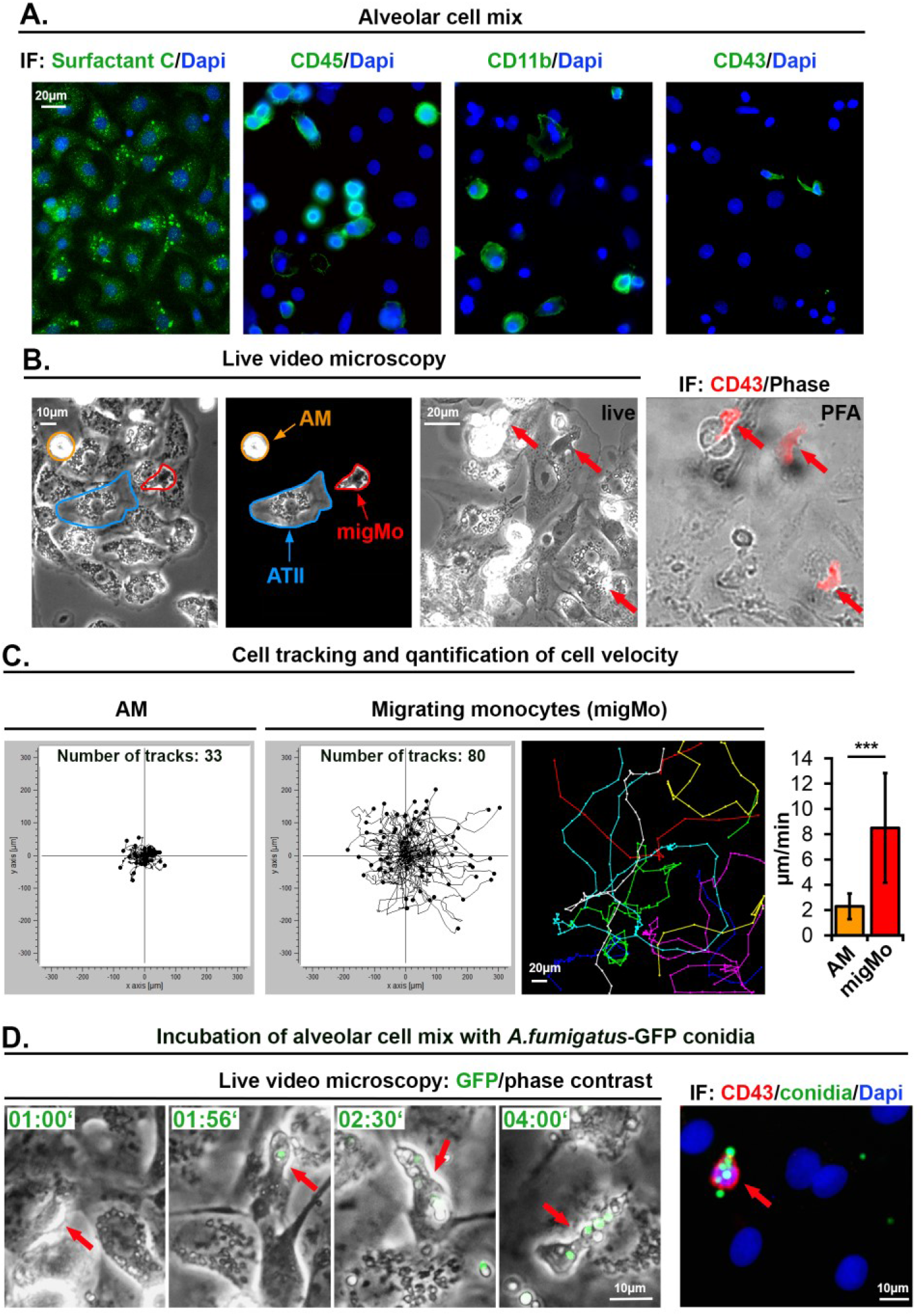
Monocytes isolated from rat alveoli migrate on top of ATII cells, internalize and transport *A.fumigatus* conidia. A. Primaiy ATII cells express Surfactant Protein C. Alveolar cell mix also contains *CD45+, CD11b*+, and *CD43*+ cells. B. Phase contrast live video microscopy revealed round slowly migrating cells (AM, orange arrow and encirculations) and fast migrating monocytes (migMo, red arrows and encirculations). Non-migrating ATII cells are marked in blue. The “live” image was taken during adding fixative immediately after the end of live video recording, red arrows point on fast migrating cells with patrolling behaviour. “PFA” is phase contrast merged with CD43 antibody in red. C. Tracks of all lung migrating monocytes observed within 6 hours, plotted in XY coordinates. Coloured image shows example of individual tracks in separate colours. Graph depicts quantification of migration velocity of AM (n=33) and MigMo (n=80). Data is shown as mean+/- standard deviation. D. Monocytes (phase contrast, red arrows) bind and internalize multiple *A.fumigatus-GFP* conidia (in green). Time is indicated as “hours:minutes”. The most right image depicts CD43 staining (red), GFP-conidia (green) and Dapi (blue). AM- alveolar macrophages, ATII – alveolar type II cells, migMo-migrating monocytes. See also supplementary **Video 1.**

Next, we performed live video microscopy in order to investigate the motility of immune cells on the surface of adherent ATII cells. We have identified two types of cellular motility: slowly moving round cells that may be AMs (orange arrows in **Figure 1B** and supplementary **Video 1**) and relatively small cells that rapidly moved over the alveolar surface. Due to the small cell size and characteristic “patrolling” migratory phenotype, the latter might be *CD43*+ non-classical monocytes (red arrows in **Figure 1B** and **Video 1**). We quantified all cells in the field of view and additionally tracked and quantified the slow and fast migrating cells. Results from three independent experiments showed a significant difference in migration velocity (2.30 ± 1.01 versus 8.5 ± 4.34 μm/min, graph in **Figure 1C**). Interestingly, when we seeded the alveolar cell isolate at 50-60% density, cells with patrolling behaviour migrated readily on the surface of ATII cells but avoided migration on empty surfaces devoid of cells (unpublished observations).

In order to further characterize the fast cells that we observed during live cell imaging, we seeded alveolar cell isolates on gridded iBidi dishes. At the end of the live imaging sequence, cells were immediately fixed using warm paraformaldehyde without perturbing the culture to enable position identification of those cells. After staining cells with anti-rat-CD43 antibodies and using the grid reference position, we identified the fast migrating cells as *CD43*+ non-classical monocytes (**Figure 1B**). Staining of these cells for rat CD4, CD103 and MPO was negative (data not shown). We have observed that nearly 50% of “live” detected fast migrating cells were lost during the fixation process, or cells did move so fast, that we could not completely overlay grid position before and after the fixation process. It might indicate transient/week adhesion of *CD43*+ monocytes to alveolar cells, which is important for fast monocyte translocation. This result also explains the difference between the percentage of cells with patrolling behaviour in live videos that was up to 9% (mean +/- 1.75%, results of 5 independent experiments) but the smaller percentage of *CD43*+ cells after fixation (4.2% +/- 1.2%, results of 3 independent experiments).

### Migrating lung monocytes internalize and transport *A.fumigatus* conidia

To further evaluate the role of migrating lung monocytes in pathogen response we challenged the primary alveolar cell isolate with 10^5^ cfu/ml of swollen conidia of GFP expressing *A.fumigatus* ^40^ Live cell imaging experiments showed that migrating monocytes bound and internalized *A.fumigatus* conidia and further transported them over the surface of ATII cells (**Figure 1D, Video 1**). We neither observed conidia germination nor fungal hyphae growth in live videos performed over 8 hours, suggesting that the presence of lung monocytes, alveolar macrophages or/and other MNPs efficiently inhibited *A.fumigatus* germination.

### Isolated patrolling monocytes inhibit *A.fumigatus* conidia growth

In order to investigate a role of migrating lung monocytes in fungal defence and to exclude the influence of other immune or ATII cells, we used magnetic cell sorting (MACS). We isolated *CD43*+ cells using CD43-beads and stained isolated cells with CD43 antibody (**Figure 2A**). From a total of 1.8×10^7^ cells in the alveolar cell isolate, CD43 selection resulted in 5.4×10^5^ *CD43*+ cells (3% of all cells in the mix).

**Figure 2.**
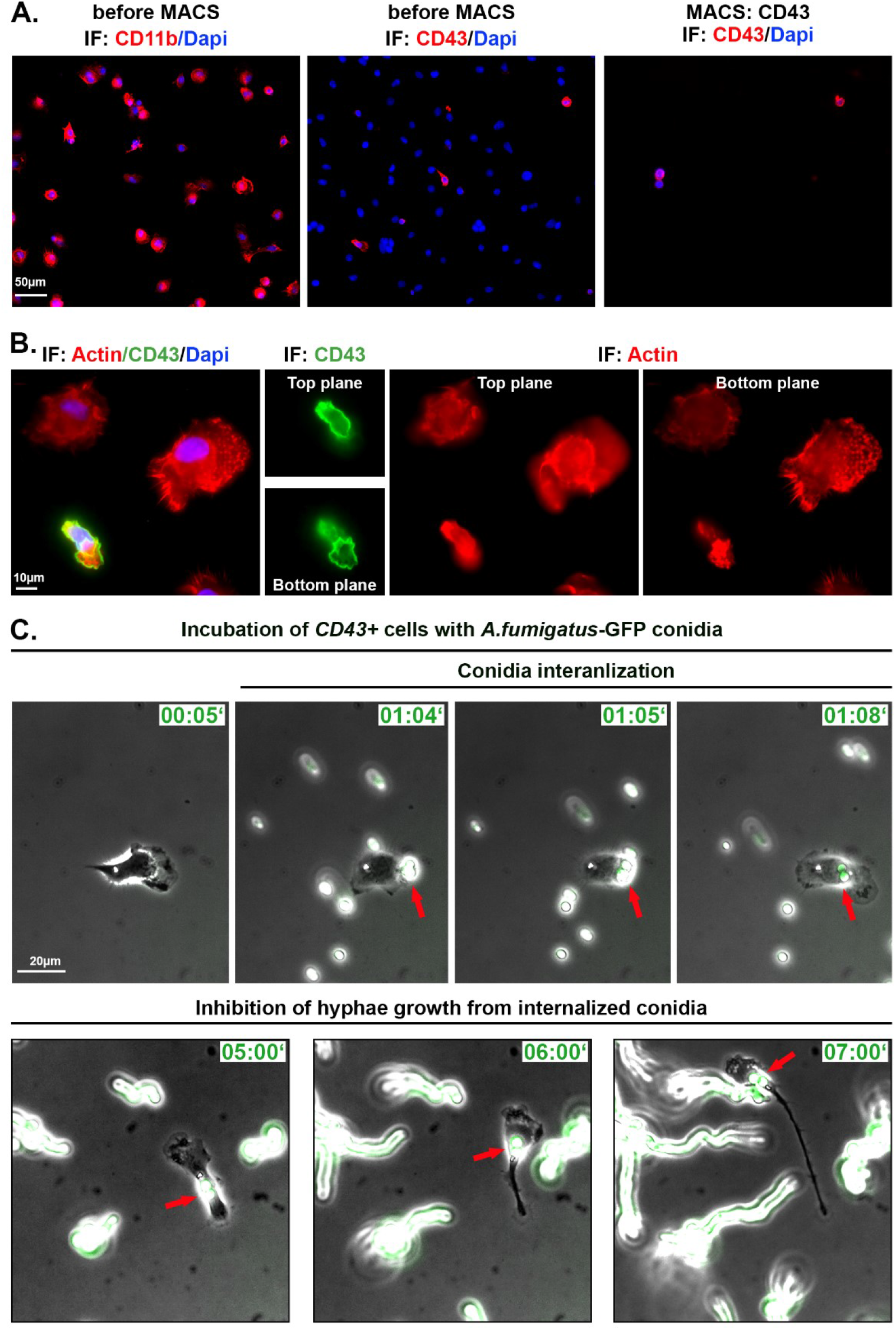
Isolated patrolling monocytes inhibit *A.fumigatus* conidia growth. A. Cells before and after MACS. IF: CD43, CD11b in red, Dapi in blue. B. Cell morphology of immune cells plated on IgG before MACS. Two focal planes are depicted. IF: phalloidin staining in red, CD43 in green, Dapi in blue. C. Isolated CD43+ cells infected with swollen A.fumigatus-GFP conidia (green). Shown are phase contrast images merged with GFP channel. Red arrows point on conidia binding and internalization into *CD43*+ cell. Time is indicated as “hours:minutes”. See also supplementary Video 2.

Isolated *CD43*+ cells were plated on IgG-coated glass cover slips/iBidi dishes, since these cells did not efficiently adhere to uncoated surface or other tested coatings, including laminin, gelatin, fibronectin, poly-L-lysine and the combination of poly-L-lysine and fibronectin (data not shown). By live cell microscopy we could observe migrating *CD43*+ cells, but in the absence of ATII cells or/and due to the IgG coating, their velocity was significantly decreased (4.59 +/- 2.43 versus 8.50 +/- 4.34 μm/min). Small cell size and polarized cell shape of *CD43*+ monocytes was clearly different from other lung immune cells as visualized by actin staining (**Figure 2B)**.

Next, we infected isolated *CD43*+ cells with 10^5^ cfu/ml of swollen *A.fumigatus-GFP* conidia. Live cell imaging experiments showed that migrating *CD43*+ monocytes internalized and transported internalized *A.fumigatus* conidia similarly to experiments in **Figures 1** and supplemented **Video 1**. We also observed non-internalized conidia that initially formed hyphae, elongated and further efficiently grew during live microscopy video (**Figure 2C**). In comparison to non-internalized conidia, internalized ones did not form hyphae and did not grow during the experiential time (red arrows in **Figure 2B, Video 2**). We also did not observe a clear conidia lysis, as the GFP-signal of internalized conidia remained essentially unchanged.

## Discussion

Migrating monocytes were recently discovered as a continuously patrolling monocytic subset in the vascular wall of blood vessels ^18,41^. “Patrolling” locomotion was first described in non-classical monocytes in the microvasculature of the dermis in mice. It was shown to be independent of the direction of the blood flow, and required integrins and LFA-1 interactions with ICAM1^41^. Carlin et al. proposed the enrichment of non-classical monocytes within capillaries and further suggested that these cells may crawl along the endothelial surfaces, phagocytose injured endothelial cells, remove debris and finally recruit neutrophils to the site of injury ^26^. Other studies characterized patrolling capacity with fast locomotion and transmigration into neighbouring tissue, thus non-classical monocytes are not restricted to the vasculature ^25,41^. In our study, we identified non-classical monocytes in the primary alveolar cell mix and show for the first time, that these cells migrate fast on the surface of alveolar ATII cells, crawling, on average, with a speed of 8.5 μm/min under cell culture conditions. Our experiments further suggest that non-classical monocytes present in the alveoli acquire precise and highly specialized molecular mechanisms to allow this fast patrolling locomotion and interaction within the alveolar epithelium. Likewise, isolated patrolling cells adhered to IgG-coated surfaces, whereas solely a small fraction of cells adhered to other coatings/uncoated surfaces. More specifically, our results have implications for a fast locomotion of patrolling cells, which is defined by the interaction with distinct receptors on the membrane of alveolar cells. This phenomenon relies on certain molecular integrin pattern expressed by patrolling monocyte when migrating in alveoli. Our preliminary data showed expression of Itgal (integrin alpha L chain), Itgam (CD11b), Itgb1 (CD29) and Itgb2 (CD18) by CD43+ monocytes, with the later to have the highest relative expression level (unpublished observation). Molecular mechanisms of patrolling cells interaction with the membrane of alveolar cells remain to be investigated.

Monocytes are leucocytes and upon infection, their cell number massively increases, accompanied by infiltration of the inflamed tissue without loss of their monocytic character ^2,12^. Tissue monocytes were shown to function in homeostatic tissue surveillance by capturing and transporting antigen to lymphoid organs ^12^ and serve specific effector functions during infection ^42^. A standardized nomenclature and definition of tissue monocytes is complex, since these cells are highly heterogeneous and can thus be described either by their function, by their gene expression profile or by phenotypical markers, and all of which can vary. Function, however, i.e. providing a critical sensing capacity with a fast locomotion ^25,41^, is key for the definition of monocytes, irrespective of their progeny. Of note, migrating tissue monocytes, positive for CD43 ^43,44^, CD14 and/or CD16 expression, can be easily separated from DCs by distinct phenotypical markers, shape and size ^12,45,46^. Furthermore, monocytes significantly differ in their more rapid turnover from DCs ^47,48^.

Tissue monocytes have also been proposed to enter and survey the lung at steady state without differentiation into DCs or macrophages ^12^. Murine monocytes have been shown to differ from other MNPs in terms of localization and trafficking: tissue-resident monocytes patrolled blood vessels and airways, whereas tissue-resident MNPs solely surveyed the latter. Regional segregation of monocytes and DCs play a crucial role, also regarding antigen uptake. This capacity appears to be differently expressed when, in a murine setting, fluorescent beads were applied intravenously or through the airway. In both cases, monocytes were more efficient in bead uptake followed by phagocytosis in comparison to DCs ^13^. This observation may point to a specific local environmental interaction of lung capillaries and –alveoli with monocytes. We observed that migrating monocytes are involved in local immune surveillance in alveoli by actively patrolling the alveolar epithelium. Rather than being recruited from the central circulation, we consider the observed cells as “resident tissue monocytes” present in the lung. This assumption is supported by the following pieces of evidence: (a) tissue monocytes can enter and survey the lung without differentiation into DCs or macrophages ^12,25,41^; b) tissue monocytes in lung differ from other MNPs in their localization and trafficking; tissue-resident MNPs only survey the airways, whereas tissue-resident monocytes survey both, blood vessels and airways ^13^. To our knowledge, we are the first group that use a primary alveolar cell mix containing alveolar cells and MNPs as well as the first ones who isolated primary migrating monocytes from the distal lung tissue.

The question concerning the functional differences between blood derived- (recruited) and lung specific (resident) monocytes remains unsolved, particularly for migrating lung monocytes. A significant amount of data was obtained on the process of monocyte transmigration from blood circulation into the lung. In contrast, only a few recent studies pointed on the importance of monocyte localization in lung compartments for immune surveillance and pathogen phagocytosis. Due to the fast locomotion and the ability to internalize pathogens (e.g. fungal conidia), these cells may represent an essential element in the alveolar defence mechanisms against pathogens. Our data suggest that by infecting the primary alveolar cell mix with fungal conidia of *A. fumigatus* patrolling cells can indeed efficiently internalize multiple living *A.fumigatus* conidia.

In described here experiments, we have observed inhibition of fungal growth inside migrating monocytes. However, we did not resolve the fate of internalized conidia. Use of recently developed fluorescent Aspergillus reporter stain (FLARE)^33^ may help to distinguish live and dead conidia. Still, in experiments with MACS-isolated *CD43*+ cells we excluded effects of other cell types present in lung tissue such as ATII cells, AMs, DCs, natural killers and neutrophils thus eliminating cell-cell interactions and impact of released pro-inflammatory mediators.

The plasticity of monocytes to respond to pathogenic threats means that different anatomical sites are likely to induce specific phenotypes. The local interaction with tissue specific cells, e.g. lung alveolar cells, as well as the regulation of above mentioned integrin cluster remains enigmatic. Further investigations may re-evaluate the role of tissue specific monocytes in health and disease, and to deepen our knowledge of monocyte patrolling during the course of pulmonary infections.

## Materials and Methods

### Reagents

DMEM medium, fetal bovine serum (FBS), L-glutamine, penicillin/streptomycin and PBS were from Capricorn Scientific, EDTA and paraformaldehyde from Merck, IgG-solution, Triton X, Tween 20, Sabouraud and BSA from Sigma. Goat serum was from Vector.

### Isolation of primary rat alveolar cell mix

The isolation of alveolar cells was carried out as previously described ^37,49^ with slight modifications. Briefly, Sprague-Dawley rats were anaesthetised, heparinised and bled out. The lungs were cleared of blood by perfusion and removed from the thorax. After lavage, the lungs were incubated with elastase and trypsin and were finally minced in DNase-solution. After adding FBS to stop the enzymatic reaction, cells were filtered through gaze (2 and 4 layers) and nylon mesh (150μm, 20μm and 7μm) and centrifuged at 130 g for 8 minutes at 4°C. For live imaging and immunofluorescence experiments, pelleted cells were re-suspended in DMEM, plated onto 10 cm plastic dished coated with IgG-solution and incubated at 37°C in a CO2-incubator for 10 minutes to remove excessive immune cells. Non-adherent cells were harvested by collecting the supernatant from the petri dishes and washing the cells at 130x g for 8 minutes at 4°C. The cells were finally re-suspended and cultured in DMEM supplemented with 10% FBS, 2 mM L-glutamine and penicillin/streptomycin at 37 °C with 5% CO2. For magnetic cell separation (MACS) experiments, cells were not plated on IgG-coated dishes, thus increasing number of MNPs in the cellular isolate.

### Magnetic Cell Sorting (MACS)

MACS was performed using MiniMACS separator and MACS columns (Miltenyi) according to the manufacture instructions. Briefly ATII alveolar cell mix was washed in MACS buffer (PBS, 2mM EDTA, 0.5% BSA), cells were centrifugated at 300xg for 10 minutes at 4°C, cell pellet was resuspended in 80μl MACS buffer and 20μl of CD11b/c MicroBeads, (Miltenyi, rat-specific), incubated for 15minutes. Further 2ml of MACS buffer was added following 300xg centrifugation at 4°C for 10 minutes. Cell pellet was resuspended in 500μl of MACS buffer, *CD43*+ cells were separated using columns on ice, washed 3 times in MACS buffer and resuspended in DMEM. Cells were seeded on IgG-coated (IgG resolved in Tris-HCl, pH 9.4, incubated for 4 hours at room temperature) iBidi dishes (y-Dish 35mm, iBidi) or on glass cover-slips and maintained in growth DMEM. For infection with *A. fumigatus* the media was replaced with antibiotic free- DMEM.

### Cultivation and growth of *Aspergillus fumigatus*

The *A.fumigatus* strain expressing green fluorescent protein (GFP) ^40^ was grown on Sabouraud 4 % glucose agar (15 g/l agar, 40 g/l D(+)-glucose, 10 g/l peptone) plates at 37 °C for 7 days until fully maturation of spores was observed. Conidia were harvested by rinsing the culture surface with sterile distil water containing 0.01% Tween20. Conidia suspension was washed once in PBS, twice in DMEM without antibiotics and counted using a haemocytometer. Freshly harvested conidia (10^6^ cfu/ml) were kept in DWEM in shaking incubator at 37°C, 160 rpm for 2 hrs to obtain swollen conidia and used in all experiments. For experiments, conidia at concentration 10^5^ cfu/ml were used.

### Immunofluorescence

Cells were fixed in 4% paraformaldehyde for 20 minutes at room temperature and washed 3 times in PBS. Cells were permeabilized with 0.05% Triton X in PBS for 10 minutes followed by washing in PBS. Blocking was performed in 10% BSA and 5% goat serum for 30 minutes. First and secondary antibodies were diluted in PBS containing 1% BSA and 5% goat serum. First antibodies were incubated for 1h followed by washing in PBS. The following antibodies were used: mouse anti-rat CD43 antibody (BioRad), rabbit polyclonal Surfactant C (Proteintech), mouse anti-rat CD11b (BioRad), mouse anti-rat CD45 (BioRad). Secondary antibodies were incubated for 1h followed by washing in PBS. The following secondary antibodies were used (all from Invitrogen): goat anti-mouse Alexa-FluorPlus488, goat anti-rabbit Alexa-FluorPlus488, goat anti-mouse Alexa- FluorPlus555. DAPI (Sigma) was used for nuclei staining. Pictures were taken using an Oxion Inverso Microscope equipped with precooled CCD camera (both from Euromex) and the ImageFocus 4, version 2.8. Software. Alternatively, images were taken using inverse Zeiss Axiovert 200M (Zeiss) equipped with a CCD camera (CoolSNAP HQ2; Photometrics), and acquisition was controlled by the AxioVision release 4.5 SP1 software (Carl Zeiss). Live video microscopy was performed using the same Zeiss setup at 37°C and 5 % CO2 conditions. Videos and cell tracking were prepared using Fiji free software^50^, Manual Tracking plug-in (Fabrice Cordeli, Institute Curie, Orsay, France) and Chemotaxis and Migration Tool plug-in (Ibidi, Martinsried, Germany). Figures were prepared in Fiji and Photoshop.

### Statistics

For quantification of migration velocity, values were transferred from Fiji into Excel program and analysed by Student’s t-test and one-way ANOVA. Results are expressed as means ± standard deviation (SD) in absolute numbers (μm/min), P ≤ 0.001 was considered significant.

### Ethical issues

Alveolar Type II cells are prepared from anesthetized and analgised Sprague-Dawley rats in the laboratory of Thomas Haller. All animal experiments and all steps of cell preparation were approved by the Institutional Animal Care and Use Committee at MUI and were conducted in conformity with the Austrian rules for animal care and testing (a license from the Austrian Government, appl. no. BMWFW-66.011/0160-WF/V/3b/2016).

## Acknowledgments

This research project was supported by the Austrian Research Promotion Agency (FFG), Grant # 856320 to Susanne Perkhofer. GFP-expressing *A. fumigatus* strain FGSC A1258 gGFP was from Genetics Stock Center, Manhattan, Kansas, USA. We thank Stephanie Angerer and Christoph Zulmin for the technical support and Sandra Schaffenrath for English corrections. We are very grateful to Irina Öttl (†2020) who performed isolation of primary rat alveolar cells.

## Supplementary video files

**Video 1 (for Figure 1B, D)** shows migration of lung monocytes on top of alveolar cells before (left video, phase contrast) and after (right video, phase contrast merged with GFP-channel) the internalization of *GFP-A.fumigatus* conidia. Orange arrows indicate AMs, blue arrows direct to ATII cells, red arrows point on migrating monocyte (migMo), green arrows indicate conidia. Time is indicated in hours:minutes. Green font on the right video indicates time after *GFP-A.fumigatus* conidia were added.

**Video 2 (for Figure 2C)** shows internalization of *A.fumigatus* conidia into MACS-isolated *CD43*+ monocyte and inhibition of fungal growth. Phase contrast merged with GFP-channel (left), GFP channel (right). Red arrows point on internalized *GFP-A.fumigatus* conidia. Time is indicated in hours:minutes.

